# Phenotype-driven parallel embedding for microbiome multi-omic data integration

**DOI:** 10.64898/2025.12.05.692119

**Authors:** Tal Bamberger, DAP Consortium, Elhanan Borenstein

**Affiliations:** Gray Faculty of Medical & Health Sciences, Tel Aviv University, Tel Aviv, Israel; Blavatnik School of Computer Science and AI, Tel Aviv University, Tel Aviv, Israel; Santa Fe Institute, Santa Fe, NM, USA

## Abstract

The human microbiome is widely recognized as a key determinant of health and disease, yet most reported links between observed microbial features and clinical outcomes remain descriptive and lack an integrated system-level perspective. Multi-omic studies of the microbiome, which jointly profile and analyze multiple molecular aspects of the microbiome via metagenomics, metabolomics, proteomics, and transcriptomics assays, offers a more comprehensive view of this system, with the potential to uncover how microbial communities and functions influence host physiology. However, integration of such multi-omic data remains challenging due to high dimensionality, major differences in data properties across omics, and the need to utilize and preserve omic-specific information. Embedding omic data in low-dimensional spaces offer a promising avenue to capture complex patterns, reduce noise, and improve downstream analysis, yet most embedding-based microbiome studies to date exhibited limited predictive power or relied on a single joint embedding of all omics thus failing to preserve omic-species properties.

To address this, we introduce PAPRICA (Phenotype-Aware Parallel Representation for Integrative omiC Analysis), an encoder-decoder framework for microbiome multi-omic integration that embeds each omic into its own latent space while jointly modeling their relationships. The model consists of parallel autoencoders trained with a loss function that promotes three objectives: (1) accurate reconstruction of each omic, (2) alignment of samples across omics such that proximity in one latent space reflects proximity in the others, and (3) alignment with a phenotype space to capture variation associated with continuous outcomes, such as fecal calprotectin levels in IBD. The resulting models support cross-omic inference and phenotype prediction from the learned latent representations, and enables integration without collapsing data into a single space. This modeling approach thus preserves omic-specific signals while capturing phenotype-associated variation.

We compared PAPRICA to four alternative models that represent successive advances in multi-omic integration architectures. We found that across two complementary tasks, predicting one omic profile from another and predicting a continuous phenotype from an input omic profile, our parallel autoencoder approach, and particularly the PAPRICA model, demonstrated better performance across three multi-omic datasets (the Franzosa IBD cohort, Lifelines DEEP and the Dog Aging Project Precision Cohort). Combined, these findings suggest that our embedding strategy effectively captures and balances omic-specific structure, cross-omic relationships, and phenotype-relevant signals across diverse datasets, offering a flexible, scalable framework for embedding complex multi-omic microbiome data and advancing our ability to gain new insights into host-microbiome interactions.

## Introduction

The human microbiome, the collection of complex and diverse bacterial communities that the human body harbors, is tightly linked to the health of the human host^1^. The gut microbiome, especially, has been extensively studied and has been shown to influence multiple crucial host processes including development, physiology, and nutrition^2^. Moreover, imbalances in the composition of the microbiome (referred to as dysbiosis) have been linked to various disease states^3^, ranging from inflammatory bowel disease (IBD)^4^ to colorectal cancer^5^ and metabolic disorders such as type-2 diabetes^6^.

In line with the growing appreciation for the role of the microbiome in health and disease, there has been a surge in microbiome profiling efforts, aiming to link patients’ health status with the composition and activity of their microbiomes. Importantly, however, the composition of the gut microbiome varies widely: individuals can differ markedly by the relative abundance of specific taxa, or by the overall structure and diversity of the community. This biological variability making it challenging to establish a precise distinction between a normal “healthy” and a disbiotic “unhealthy” microbiome^7^, let alone to identify specific microbial components or processes that are linked to a particular disease. This variation is influenced by a multitude of factors, including the host’s diet, genetics, demographics, and environment, all of which are known to modulate microbiome composition and function. Understanding the complex associations between the microbiome and health thus requires a comprehensive approach that characterizes key components like microbial diversity, metabolic activity, and host-microbe interactions, as well as their functional roles in physiological processes.

Aiming to address this challenge and to provide a more complete, systems-based, and multi- faceted view of this complex biological system, microbiome researchers often adopt a multi-omic study design. By integrating various ‘omic’ technologies such as metagenomics, metabolomics, genomics, and other molecular profiling approaches, these studies aim to offer a comprehensive understanding of biological phenomena across multiple functional and organizational levels. Indeed, in recent years, this approach has gained significant traction in microbiome research, with an explosion of microbiome multi-omic studies that simultaneously assayed multiple aspects of community structure, including, for example, both taxonomic and functional compositions^8–13^ and more recently, both fecal microbiome and fecal metabolome^4,5,14–17^. This latter study design, combining metagenomics, which profiles microbial DNA, with metabolomics, which assays small-molecule metabolites, has allowed researchers to gain valuable insights into the functional interplay between microbial communities and their host environments. Importantly, however, despite these advances, integrating diverse multi-omic datasets remains a significant challenge due to high dimensionality and vast differences in data structure and distributions across omics, ranging from count-based matrices to continuous intensity measurements^18^. Different omics may also differ substantially in scale, with some omics yielding thousands of features and others only a few dozens, further complicating joint analysis and normalization^18^. Moreover, gaining meaningful and interpretable insights from such integrative analyses is also challenging given the distinct molecular layers that the various omics represent, each associated with a unique biological context^19^.

One promising approach to overcome some of these challenge and successfully integrate multi-omic data is using embedding methods^20–23^. This approach relies on the assumption that while microbiome omic data generally reside in a high-dimensional space, biological processes, regularities, and inter-omic dependencies (and hence meaningful biological insights) often occur in, and can be captured through lower-dimensional representations. Embedding methods thus aim to project (i.e., “embed”) high-dimensional multi-omic data into a shared lower-dimensional latent space, enabling the simultaneous analysis of multiple data types. Moreover, by capturing underlying relationships between different omic layers, embedding-based approaches facilitate identification of cross-omic dependencies and interactions that are otherwise difficult to discern. Indeed, such integrative analyses have shown promise in linking molecular variations to phenotypic traits and uncovering multi-omic functional associations^24–27^.

While many different methods to embed high-dimensional complex data have been introduced over the years, nowadays, researchers often resort to *autoencoders* to achieve this task. Briefly, an autoencoder consists of an encoder network, which compresses input data into a lower-dimensional space, and a decoder network, which reconstructs the input from this representation. By training these models on available data and minimizing reconstruction error, autoencoders uncover underlying structures and non-linear relationships within the data. As such, autoencoders form a powerful model for generating representations that capture meaningful patterns of complex data^28^, and have indeed proven effective in embedding microbiome data^29^. Moreover, the ability of autoencoders and related encoder-decoder deep learning methods to model intricate patterns makes them particularly useful for integrating diverse multi omic data, with several extensions of such frameworks supporting the incorporation of multiple omic layers^30–33^. For example, a recently proposed approach for integrating multiple modalities involved training a model on paired ECG and cardiac MRI data to map each modality into a common latent representation^34^. This mapping is guided by a contrastive loss, which enforces the constraint that paired samples are embedded closer together while unrelated samples are positioned further apart, and was shown to support downstream tasks such as phenotype prediction and clustering^35^.

In the microbiome research domain, however, the application of encoder decoder-based methods for analyzing and integrating multi omic data is a more recent development, with relatively few examples of such studies^35–38^. For example, *Deep in the Bowel*^39^ , a neural encoder-decoder network, predicts gut metabolites from microbiome data by learning a sparsely weighted latent representation of microbial abundances. This learned space captures clinically meaningful microbe-metabolite relationships and can accurately predict inflammatory bowel disease (IBD) status and treatment outcomes. Chen *et al*. introduced *CEM_HN (Combined Embedding Model based on Heterogeneous Network)*^38^, which infers microbe-metabolite interactions using three embedding components: node embedding for local interactions via a graph convolutional network; paired embedding for associations using an encoder-decoder with bridge nodes; and combined embedding for a global representation that improve prediction. Another model, *LOCATE (Latent variables Of miCrobiome And meTabolites rElations)*^40^, builds a latent representation of microbiome data to better predict metabolites and host phenotypes. *LOCATE* uses a neural network to transform microbiome data into a latent space and relates this to metabolite concentrations via a linear model, effectively predicting host phenotypic traits such as disease status and clinical outcomes. Other methods aim to integrate microbiome multi-omic data into a shared latent space while incorporating an additional loss function designed to classify samples into categories using this shared lower dimensional space. A recent study, for example, introduced the *Multimodal Variational Information Bottleneck (MVIB)*^36^, a deep learning model that integrates shotgun metagenomic sequencing data (species-relative abundances and strain-level markers) and metabolomic data to predict disease. By learning a joint representation of these data modalities, *MVIB* improves disease prediction accuracy. Cui *et al*. presented the *microbiome-based multi-view convolutional variational information bottleneck (MV-CVIB)*^37^, an approach for predicting metastatic colorectal cancer using gut microbiome data. The model employs a multi-view convolutional variational information bottleneck that integrates relative abundance data and nearest neighbor information derived from 16S rRNA sequencing to distinguish between metastatic and non-metastatic colorectal cancer. Finally, *Incomplete Multi-Omics Variational Neural Networks (IMOVNN)*^35^ enhances disease prediction and biomarker identification by learning both marginal representations of each dataset and joint latent representations. *IMOVNN* addresses challenges posed by incomplete multi-omic data and offers interpretability by selecting the most relevant features.

It is important to note, however, that despite these exciting results, existing methods face several limitations. First, while some models achieve high prediction accuracy, they often lack interpretability and fail to provide insights for directly linking latent space features to underlying biological phenomena. Additionally, many methods primarily focus on predicting categorical phenotypes, such as disease state, and have not been shown to effectively predict continuous phenotypes, such as disease severity. Many existing models are also not well-equipped to handle missing or partially-missing omic data, a frequent challenge in these complex datasets. This limitation reduces their effectiveness in real-world microbiome studies, where often some samples may lack certain omic layers, making robust integration and analysis more difficult. Finally, and perhaps most importantly, many of the above architectures are based on learning a *shared* latent representation for the various omics. While this approach allows cross-omic prediction and potentially reveals dependencies between omics, by forcing all omics into a joint representation it may obscure unique information inherent to specific omics. Combined, these challenges underscore the need for novel approaches capable of integrating a broad range of omic data, support cross-omic and continuous phenotype prediction, and rely on architectures that preserve omic-specific regularities and facilitate more intuitive and biologically informed analyses.

With that in mind, here, we present five modeling approaches designed to address key challenges in multi-omic integration and to leverage the complementary strengths of different design choices. We use these models to predict one omic layer from another (accommodating incomplete datasets), with a particular focus on further aligning learned latent spaces with biological phenotypes and applying the resulting representations for phenotype prediction. Collectively, these five models represent an evolution of multi-omic integration architectures: from simpler, previously introduced designs to more complex, novel, and flexible ones, thereby highlighting how architectural variations impact applicability, predictive performance, and interpretability.

## Results

### Encoder-decoder models for multi-omic integration

In this study, we propose and explore five autoencoders-based methods tailored for multi-omic integration. These models employ various encoder-decoder architectures designed to capture the intricate relationships between different omics datasets and between omics and phenotypes, in the hope of further improving downstream analytical tasks, such as phenotype prediction. The proposed models increase in complexity, ranging from simple encoder-decoder architectures that have been previously applied to analyze omics in microbiome research (and in other domains), to more complex and novel architectures that aim to address the challenges described above. Specifically, we focus on the following five models:

1. **The D (Diagonal) Model**: The D model employs a simple encoder-decoder architecture designed to predict one omic profile from another. The encoder embeds the first omic profile into a lower-dimensional latent space, and the decoder predicts the second omic profile from this latent representation. This architecture was used, for example, in Le *et al.*^39^ (Figure 1A).
2. **The Y Model**: The Y model utilizes an encoder-decoder architecture designed to predict one omic profile from another (as in the D model above), but do so while also reconstructing the first omic profile. This dual objective ensures that the latent space captures not only the features relevant for predicting the second omic but also those unique to the first omic (Figure 1B).
3. **The X Model**: The X model employs an encoder-decoder architecture designed to predict and reconstruct both omics, using a *shared* latent space, *with the input being **either** the first omic profile or the second omic profile*. The model is trained with alternating input (e.g., using the first omic during even epochs and the second omic during odd epochs), allowing the model to support incomplete datasets (i.e., with a certain omic is available for some samples but not for others), to leverage complementary information across the two omics (when both are available for the same set of samples), and to predict each omic from the other while learning both shared and unique features (Figure 1C).
4. **The Parallel Models**: The two “parallel models” (P_d_, and P_dp_) share a common architectural framework, consisting of two (or more) autoencoders trained in parallel, with each autoencoder corresponding to a specific omic dataset. In contrast to the models above, this approach offers the unique benefit of generating a *distinct latent space representation for each omic dataset*, thus allowing the autoencoders to preserve the distinct structures and regularities of each omic, as well as any modality-specific patterns. To this end, each autoencoder is using its own reconstruction loss function, allowing it to independently capture the characteristics of its respective omic dataset (Figure 1D). We specifically propose and study two variants of this architecture, which differ primarily in the loss function components used for coupling the various omics:

- Parallel Model with Distance-based Coupling (**P_d_**): The P_d_ model aims to align the structure and distribution of the different omics’ latent spaces by augmenting the loss function with an *inter*-*omic distance loss* component, ensuring that samples close to one another in one omic’s latent space are also close in the other latent spaces.
- Parallel Model with Distance-based and Phenotype Coupling (**P_dp_**): The P_dp_ model aligns the different omics using the omics distance loss component as in the P_d_ model above, but further aims to align the omic-specific latent spaces with a phenotype space, by augmenting the loss function also with a *phenotype distance loss* component. This component ensures that the latent spaces also capture distances between samples in the phenotype space, integrating available phenotypic information during training to enhance the biological relevance of the latent representations. For future reference, we term this model PAPRICA for Phenoype Aware Parallel Representation for Integrative omiC Analysis.

**Figure 1:**
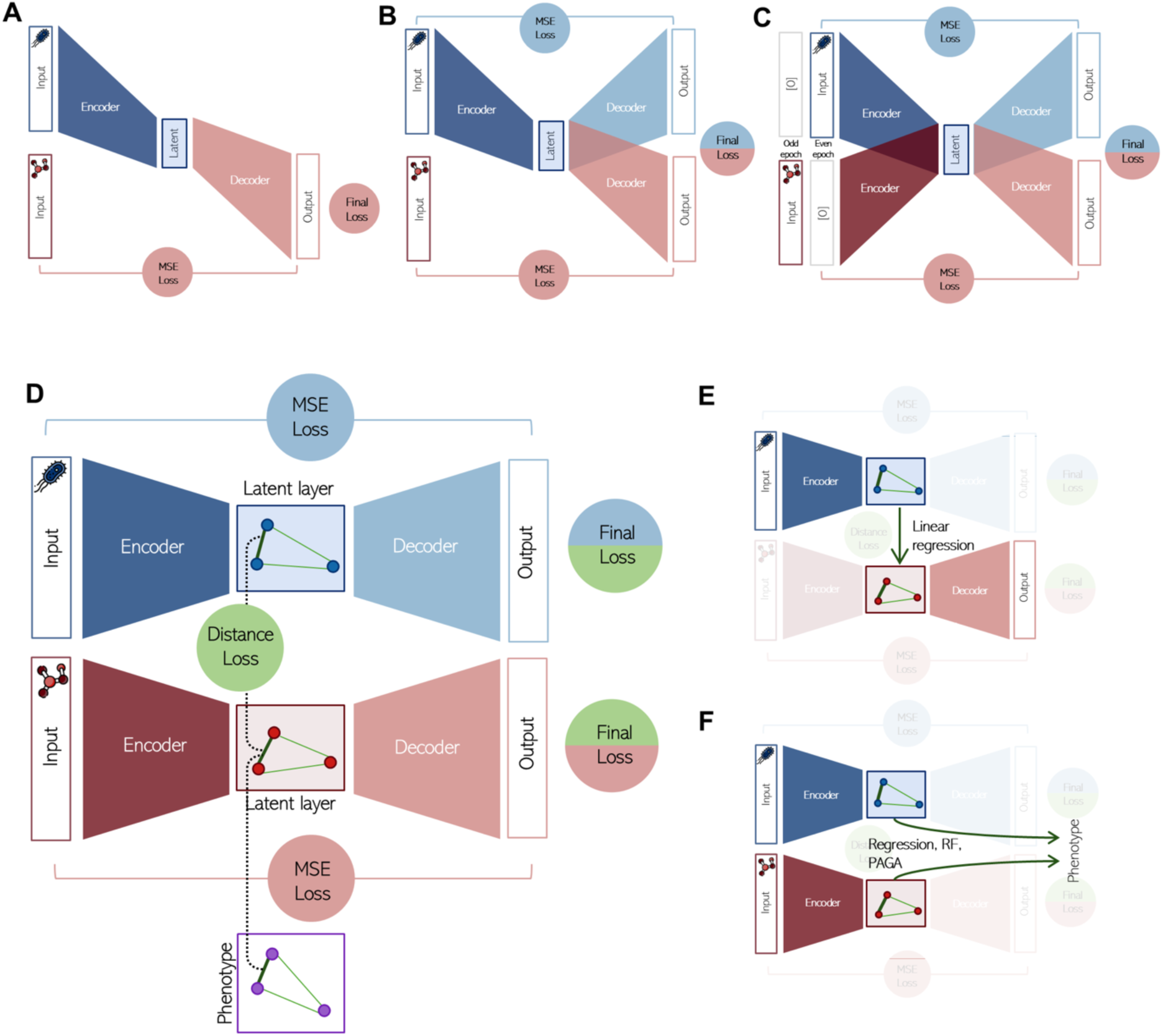
Architectures of the five proposed encoder-decoder models for multi-omic integration. **A.** The D model predicts one omic profile from another using a simple encoder-decoder architecture. **B.** The Y model predicts one omic profile from another, while also reconstructing the original omic, ensuring a comprehensive latent space. **C.** The X model alternates inputs between omics and reconstruct all omics during training to allow the model to capture complementary omic information and reconstruct all omics based on any input omic. **D.** The Parallel models incorporate multiple autoencoders to generate omic-specific latent spaces, aligning the overall structure of the latent spaces between omics using an *inter-omic distance loss* component (model P_d_), or also integrating phenotype-based alignment using a *phenotype-distance loss* component (Model P_dp_). **E.** A scheme of predicting one omic from another using the parallel models. **F.** Phenotype prediction from the latent spaces of the parallel models.

Notably, the two parallel models, P_d_, and P_dp_, use a dynamically weighted loss function that initially prioritizes accurate omic reconstruction, and as training progresses, gradually shifts to prioritize inter-omic coupling (and in the P_dp_ model, also phenotype coupling). This design allows the models to first focus on preserving omic-specific characteristics, and then further promote cross-omic and phenotype-informed alignments.

### Model assessment approach

To analyze and compare the performance of the various models, we focus on two complementary tasks. The first is cross-omic prediction, i.e., predicting one omic profile from another. Cross-omic prediction in the D, Y, and X models is inherent to the model architectures (although limited to using just one of the omics as input in the D and Y models). To support cross-omic prediction also in the parallel models, after training the autoencoders, a simple linear regression model is fit (based on the training data alone) to map the latent representation of one omic to that of another. Cross-omic prediction can then be done by using the encoder of the input omic to get a latent representation of that omic profile, mapping this representation to the latent representation of the desired output omic with the linear regression model, and then passing this predicted latent representation through the decoder of the other omic, generating a full prediction of the output omic’s features (Figure 1E). The second task is phenotype prediction, i.e., predicting a continuous phenotype from an input omic profile. As noted above, previous studies have shown that latent representations often capture important regularities of high dimensional data, and that accordingly, predicting a phenotype of interest from such latent representations (rather than from the original data) may result in a more accurate prediction. We thus tested whether the resulting latent representation of each model indeed improves phenotype prediction. It should also be noted that in contrast to the simpler models, the parallel models produce a separate latent space for each omic, thus support predicting the phenotype based on each omic independently (Figure 1F).

Finally, while all models above aim to integrate two (or more) omics without any specific assumption about the omics used, in the analyses below, we focused on integrating data from microbiome-metabolome studies, a widespread and important multi-omic study design. We specifically use initially fecal microbiome and metabolomic data from a case-control study of inflammatory bowel disease (IBD) patients^14^. In this cohort (including 152 individuals), each patient further had information about measured levels of fecal calprotectin, a marker of intestinal inflammation, which was used as the continuous phenotype of interest.

We further tested all models on additional data, using specifically the Lifelines DEEP cohort^41^, a multi-omic resource with fecal microbiome and blood metabolome profiles from 1,248 individuals, and the first-year Precision cohort of the Dog Aging Project (DAP)^42^, which includes fecal shotgun metagenomes and blood metabolomics from over 600 dogs.

### Loss curves analysis and training dynamics

To first assess training dynamics, we monitored the loss curves on *unseen test data* throughout the training process, aiming to ensure that while the models learned from the training data, their performance on unseen test data remained stable and improved as expected (Supplementary Figure 1). Notably, the different models are trained with a variety of loss functions, ranging from a loss function that is based on the prediction of a single omic (e.g., model D), to a loss function that balances the reconstruction of multiple omics with the alignment of latent spaces and phenotype (e.g., model P_dp_). We accordingly evaluated both the final loss values and their individual components for each model, focusing on the convergence behavior and the stability of the loss over time to identify potential underfitting, overfitting, or unstable optimization.

The D model, perhaps not surprisingly, exhibited steady convergence (Supplementary Figure 1A), reflecting its simplicity and straightforward predictive focus. The Y model showed slightly slower convergence, likely owing to the need to optimize two different objectives, namely reconstructing the first omic and predicting the second omic; notably, however, it ultimately reached a comparable loss value (Supplementary Figure 1B). The X model displayed a similar behavior to the Y model (Supplementary Figure 1C), with the alternating input approach indeed allowing it to successfully learn to reconstruct/predict the two omics from either the microbiome profile or metabolome profile alone (Supplementary Figure 1C, lower panels). The P_d_ and P_dp_ models demonstrated somewhat less stable loss curves compared to other models, likely due to the complexity of the loss function and the requirement to balance multiple loss components. Importantly, however, the overall dynamics still indicate a general reduction in loss values, reflecting the model’s capacity to learn effectively despite the added complexity (Supplementary Figure 1D-E). Overall, these results suggest that while the various models somewhat differed in convergence dynamics and complexity, they all successfully learned the desired objective/s from the data.

### Prediction of one omic from another

Having shown that all models exhibited successful training dynamics, we next turned to evaluate all models based on their ability to predict one omic profile based on another. As noted above, while the first three models (D, Y, X) can generate cross-omic predictions directly owing to the shared latent space, the parallel models utilize a linear regression model of map the latent space of the input omic to that of the output omic (Figure 1E; see Methods for details).

We again focused on predicting metabolomic profiles from microbiome profiles in the IBD cohort described above, and defined well-predicted metabolites as those with a Spearman’s correlation greater than 0.3 between their real and predicted values across samples, with an FDR corrected p-value < 0.1. Examining the number of well-predicted metabolites obtained by each model (across different hyperparameters) revealed an overall increase in prediction performance from the D model (which exhibited the poorest performances, with 118.45 well-predicted metabolites on average), to the Y and X models (153.23 and 167.875, respectively), and to the two parallel models (181.00 and 169.725; Figure 2A, Supplementary Table S1). This trend suggests that the addition of a microbiome reconstruction objective, as introduced in the Y and X models compared to the D model, may help stabilize the learning process and result in latent representations that better support metabolite prediction (despite the successful learning dynamics observed in model D above). Among the parallel models, the P_d_ model performed somewhat better than the P_dp_ model, potentially suggesting that the phenotype-distance loss introduces some complexity that hinders the training process, especially when the sample size is limited. Notably, however, the parallel models also appear to be somewhat more sensitive to hyperparameter variations, as indicated by the increased variance in their performance across different hyperparameter settings. As an alternative approach to quantify metabolome prediction, we also calculated the mean Spearman’s correlation between observed and predicted metabolite levels across all metabolites. Using this metric showed qualitatively similar patterns in terms of model performances, with mean correlation values of 0.269, 0.298, 0.307, 0.325, and 0.310 for the D, Y, X, P_d_, and P_dp_ models, respectively (Figure 2B).

**Figure 2:**
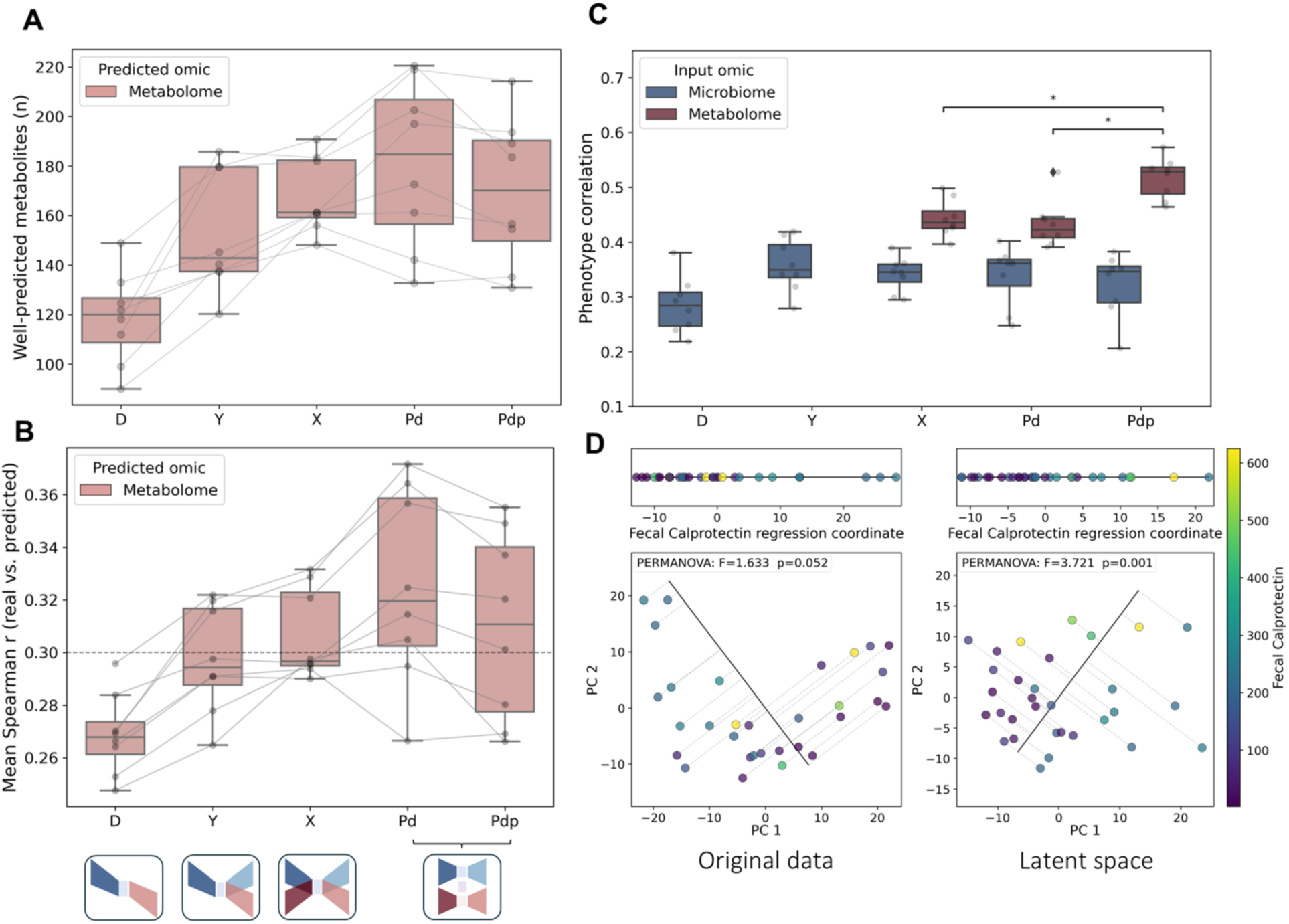
Evaluation of model performance in cross-omic and phenotype prediction. **A.** Box plot representing the number of well-predicted metabolites (mean Spearman’s r > 0.3, FDR < 0.1) for each of the five models. Each dot represents a single run under specific hyperparameters, with lines connecting runs that share the same hyperparameters across different models. **B.** Box plot showing the mean Spearman’s correlation between true and predicted metabolites for each of the five models evaluated. As in panel B, each dot represents a single run under specific hyperparameters, with lines connecting runs that share the same hyperparameters across different models**. C.** Box plot illustrating phenotype prediction performance (mean Spearman’s r between real and predicted phenotype values). The color of the boxes indicates the input omic (blue: microbiome, dark red: metabolome). Statistical significance was determined using the Mann-Whitney U test, with asterisk denoting significance after FDR correction (p<0.05). **D.** Principal Component Analysis (PCA) of the metabolome data in the unseen test samples, using either the original metabolome data (right), or the latent space representation obtained from the P_dp_ model (left). Lower panels: Scatter plots of the various samples using the first two PC and colored by fecal calprotectin level. A regression line is included to show the association between fecal calprotectin and the principal component space, with dashed lines indicating each sample’s orthogonal projection on this regression line. Upper panels: the same test samples collapsed onto the 1D regression coordinate. Association to the fecalcalprotectin was quantified via PERMANOVA with resuls reported in the panels. PCA of the train data can be seen in Supplementary Figure 5.

We also evaluated the sensitivity of the above results to the selection of initial weights of the models or to other stochastic elements in the training process by running each model with the same hyperparameters 10 times. We confirmed that the variance due to model initialization was substantially smaller than the variance across models and hyperparameters (Supplementary Figure 2). Additionally, we assessed whether well-predicted metabolites remained consistent across different models and hyperparameters. Our analysis confirmed that generally similar metabolites were well-predicted across the various models and hyperparameters, suggesting that predictive performance is largely driven by true signal in the data rather than stochastic variation in model training (Supplemetary Figure 3, Supplementary Table S2).

**Figure 3:**
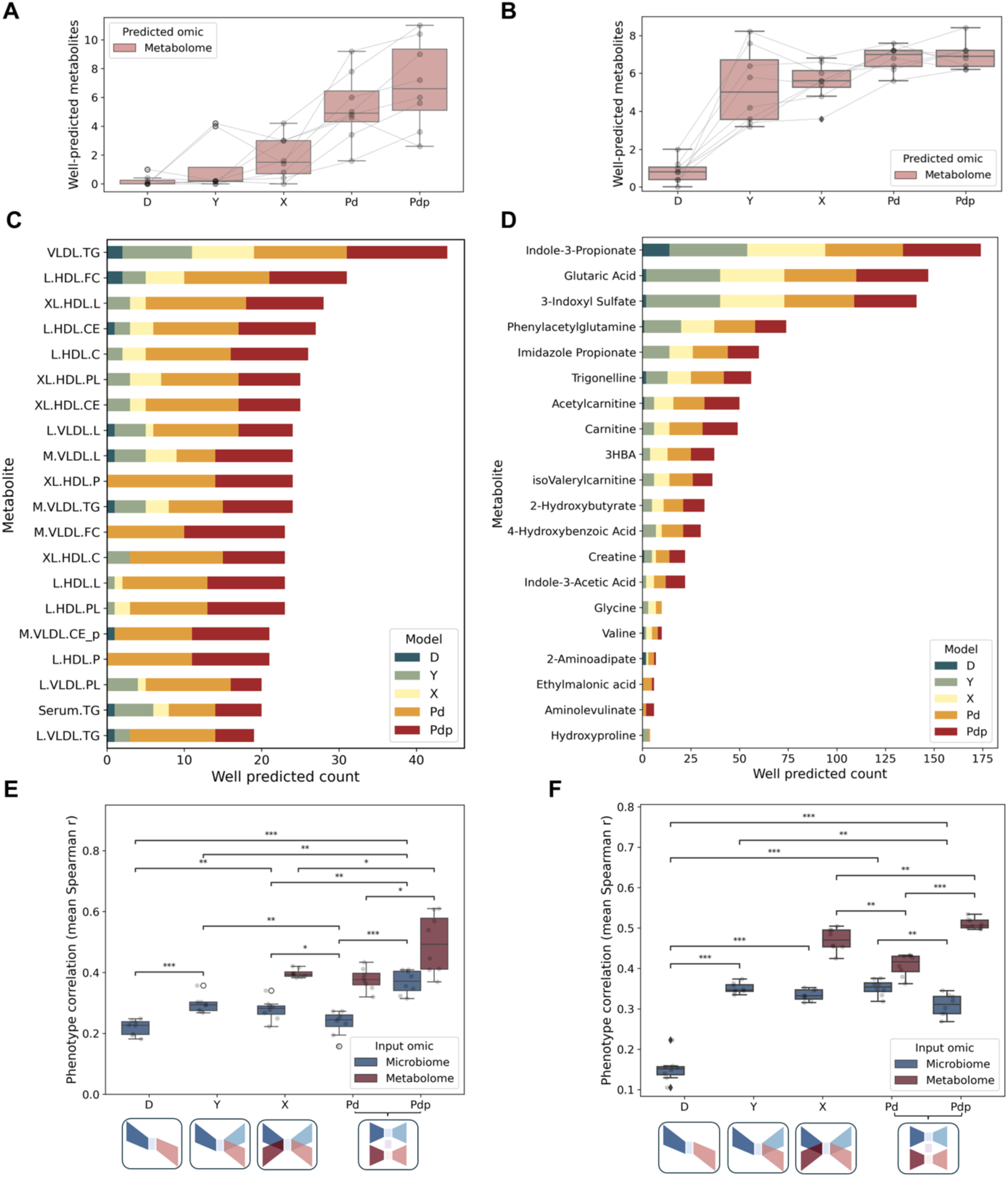
Model performance in cross-omic and phenotype prediction across the Lifelines and DAP datasets. **A-B.** Box plots showing the number of well-predicted metabolites (mean Spearman’s r > 0.3, FDR < 0.1) for each of the five models in the Lifelines (A) and in the DAP (B) datasets. See legend of Figure 2A for more information. **C-D.** Stacked bar plots illustrating the number of times each metabolite was classified as well-predicted across different hyperparameter settings and cross-validation folds in the Lifelines (C) and in the DAP (D) datasets. Only the top 20 metabolites are shown. The stacked colored segments within each bar represent different models. **E-F.** Box plots showing phenotype prediction performance from the latent space for each model (mean Spearman’s r between real and predicted phenotype values) in the Lifelines (E) and DAP (F) datasets. See legend of Figure 2C for more information.

### Phenotype prediction based on latent representations

We next turned to evaluate whether (and how well) the latent spaces generated by each model can be used predict a continuous phenotype, namely, in our dataset, fecal calprotectin levels. Phenotype prediction was done using random forest (RF), a simple and commonly-used algorithm that performs well with limited hyperparameter tuning and that can model complex, non-linear relationships effectively and robustly^43^ (using linear regression instead of random forest showed qualitatively similar behavior; Supplementary Figure 4). Notably, the D and Y models are based on a single omic encoder (microbiome in our case), and can thus be used to predict phenotype only based on input profiles from that omic. The X model, in contrast, employs a separate encoder for each omic (albeit using a shared latent space), and can accordingly be used to predict the phenotype based on any omic input. In our case, we thus evaluated its ability to predict the phenotype based on either microbiome or metabolome inputs. Finally, as noted above, the parallel models employ a different autoencoder *and* produce a separate latent space for each omic, thus allowing us to evaluate their predictive capacity using either microbiome or metabolome inputs (Figure 1F). This feature of the parallel models (and to some extent, also of the X model), supporting multiple input types for predicting a phenotype of interest, is a significant advantage for these models, providing complementary perspectives on phenotype association while still integrating multiple omics.

Indeed, this enhanced and flexible predictive capacity of the more complex models proves beneficial when assessing the predictive power of the various models (Figure 2C, Supplementary Table S3). Specifically, when using microbiome profiles as input, all models exhibit generally similar performances in predicting the phenotype (mean Spearman’s r of 0.285, 0.358, 0.341, 0.340, and 0.322 for the D, Y, X, P_d_, and P_dp_ models, respectively), with the simple D model being the least successful. These poor performances of the D model may highlight the advantage of using latent representations that capture both shared and unique features of the various omics. More importantly, however, when using metabolomic profiles as input (available only in the X, P_d_, and P_dp_ models), the average correlation between observed and predicted fecal calprotectin increased across all these models (mean Spearman’s r: 0.443, 0.433, and 0.517 for the X, P_d_, and P_dp_ models, respectively). The P_dp_ model, specifically, achieved significantly higher correlations than the other models when metabolomics data were used as input (Mann-Whitney U, FDR < 0.1), reflecting stronger alignment of the latent space with the fecal calprotectin in both training and test datasets. Following up on these findings, we next examined whether the latent space constructed by the Pdp model better captures fecal calprotectin related variation than the original feature space. Comparing the association of fecal calprotectin with these space , we indeed found a stronger association in the latent space representation vs the original space (F = 3.721, p = 0.001 vs F = 1.633, p = 0.052, PERMANOVA test), further suggesting that incorporating phenotype information during training guides the latent space toward clinically relevant variation (Figure 2D, Supplementary Figure 5).

### Evaluating the models’ performances across additional datasets

To confirm the validity and robustness of our findings above, we finally evaluated the performances of the various models using two additional and independent larger datasets. The first was obtained from the Lifelines DEEP cohort^41^, a multi-omic dataset with fecal microbiome and blood metabolome data from 1,248 individuals, complemented by additional biological and phenotypic data. The second dataset was obtained from the first-year Precision cohort of the Dog Aging Project (DAP)^42^and includes fecal shotgun microbiome sequencing and blood metabolomics from over 600 dogs. In both datasets, we used the chronological age as the continuous phenotype to be predicted.

As above, we first examined the loss dynamics (when applied to unseen test data) during the training of the various models on these datasets. Overall, observed trends were consistent with those seen in the smaller dataset above, although here, the larger sample size resulted in a more stable behavior (Supplementary Figure 6-7). We then turned to examine the models’ ability to predict metabolites levels based on microbiome data. It is important to note, however, that in contrast to the dataset analyzed above, metabolome data in both Lifelines DEEP and DAP were derived from blood rather than fecal samples. This is clearly expected to markedly weaken the metabolome’s association with the microbiome, and in turn to result in substantially fewer well-predicted metabolites^44,45^. Consistent with this assumption, a Mantel test of the correlation between the microbiome and the metabolome distance matrices showed a strong global association in the prior stool-stool dataset (Mantel r = 0.226, p = 0.001), but substantially weaker associations in Lifelines DEEP (r = 0.044, p = 0.051) and in DAP (r = 0.066, p = 0.001). Indeed, in the Lifelines DEEP dataset, the mean number of well-predicted metabolites was only 0.20, 1.13 and 1.80 for the D, Y, and X models, respectively, with the parallel models again exhibiting better performances (5.30, and 6.93 for the P_d_, and P_dp_ models, respectively; Figure 3A, Supplementary Table S1). This again underscores the parallel models’ ability to integrate microbiome and metabolome data more effectively, even when the association between these two omics is weaker. Similar patterns were also observed in the DAP dataset, with the parallel models outperforming all single-network models (mean number of predicted metabolites 0.83, 5.30, 5.60, 6.76, and 6.93 for the D, Y, X, P_d_, and P_dp_ models, respectively; Figure 3B, Supplementary Table S1). We further evaluated the robustness of metabolite predictions across multiple runs, and overall consistency in metabolite prediction across models and runs. This analysis confirmed that in both datasets metabolites generally exhibited similar predictability across different models and hyperparameter settings, indicating consistent model behavior (Figure 3C-D, Supplementary Figures 8-9, Supplementary Tables S4-S5). For example, indole-3-propionate, glutaric acid, and 3-indoxyl sulfate, were all well predicted in the DAP dataset across runs and models. These metabolites are known to originate from gut microbial pathways, especially bacterial tryptophan catabolism, and to shape host physiology by supporting barrier integrity, modulating inflammation, and engaging energy metabolism^46^. In particular, indole-3-propionate is gut-protective^47^, 3-indoxyl sulfate has been proposed as a dysbiosis marker^48^, and glutaric acid correlates with host energy-metabolism gene expression^49^. We finally evaluated the ability of the each model to predict a continuous phenotype, namely age, in these two datasets (Figure 3E-F, Supplementary Table S3). As above, microbiome based prediction was generally worse than metabolome based prediction (with the D model performing especially poorly), and the P_dp_ model achieving the best prediction accuracy (using metabolome input for the DAP dataset and either microbiome or metabolome input for the Lifelines dataset). Combined, these findings suggest that our observations above concerning the predictive capacity of the various models are broadly consistent across datasets and input modalities, though effect sizes may vary by cohort.

## Methods

### Datasets description

We analyzed data from three independent resources, each comprising matched shotgun metagenomic and metabolome profiles.

We used fecal microbiome and fecal metabolome data from a previously published case-control study of 220 individuals by Franzosa et al.^14^, including 88 individuals diagnosed with Crohn’s Disease (CD), 76 with Ulcerative Colitis (UC), and 56 healthy controls. Fecal calprotectin level, a potential marker for inflammation and disease activity, was available for 152 patients (CD: 65, UC: 46, and healthy controls: 45). To address outlier values, samples with fecal calprotectin levels exceeding 2,000 were replaced with the maximum level observed among the other samples (625).

We used fecal microbioms and serum metabolites from the Lifelines cohort^50^. Lifelines is a multi-disciplinary prospective population-based cohort study examining in a unique three-generation design the health and health-related behaviours of 167,729 persons living in the North of the Netherlands. It employs a broad range of investigative procedures in assessing the biomedical, socio-demographic, behavioural, physical and psychological factors which contribute to the health and disease of the general population, with a special focus on multi-morbidity and complex genetics. From the Lifelines Deep cohort^41^, a subset of the extensive population-based Lifelines study, we used microbiome and metabolomic data from 990 individuals. In our analysis, we utilized the age as the continuous phenotypes.

For the Dog Aging Project (DAP) Precision Cohort^51^, a large-scale initiative investigating how genetics, lifestyle, and environmental factors contribute to aging in companion dogs^42^, we analyzed data of 618 dogs from the Precision Cohort. This cohort includes paired fecal microbiome and blood metabolome profiles, along with detailed phenotypic information. As in the Lifelines Deep cohorts, we used age as a continuous phenotype.

### Microbiome data processing

Across all datasets, metagenomic data were processed using a consistent pipeline: Shotgun metagenomic sequencing data was processed using fastp^52^ for quality control, Bowtie2^53^ for host read filtering, and Kraken2-Bracken^54,55^ for taxonomy assignments, with the Genome Taxonomy Database (GTDB)^56^ serving as the reference. Abundance values were converted to relative abundances summing to 1 in each sample. Low-abundance taxa were filtered based on dataset-specific thresholds; Franzosa: mean > 0.001 & prevalence > 2%; Lifelines DEEP: relative abundance > 1% in ≥1% of samples; DAP: mean > 0.001 & prevalence > 20%). Finally a small constant (1e-9) was added to all species abundance values before applying a Centered Log-Ratio (CLR) transformation to address compositional data challenges. For Franzosa et al., we used the published species relative abundance table from the gut microbiome-metabolome dataset^57^, processed using the same pipeline, retaining 188 species. For Lifelines DEEP, shotgun data were downloaded from the European Genome-phenome Archive (EGA) using the pyEGA3 toolkit (v5.2.0) and processed as above, retaining 213 species. For DAP, raw reads were processed as above, resulting in 368 species.

### Metabolome data processing

Metabolome preprocessing varied by dataset, based on platform and data availability. Specifically, in the Franzosa et al. dataset, we downloaded from the original study the processed LC-MS metabolome data and metabolite-to-class conversion table. Features present in only 50% or less of the samples were filtered out, resulting in 7,111 metabolites out of 8,848. Zero values were imputed by replacing them with half of the minimum non-zero value for each metabolite. Metabolite abundances were then aggregated at the class level by summing the abundances of all metabolites within each class, yielding 454 class-level metabolites. To reduce redundancy, highly correlated features (Spearman r > 0.8) were removed, resulting in 398 metabolites classes used for analysis. Finally, a Centered Log-Ratio (CLR) transformation was applied. In the Lifelines DEEP cohort (which was assayed using the nuclear MR method^58^), we applied the same filtering and imputation steps as above, and a small constant (1e-9) was added before applying CLR transformation to the 232 retained metabolites. For DAP, we used available processed LC-MS metabolome data, including 137 metabolites, as described in Harrison et al.^59^.

### Notation and definitions

Although the various models can accept an arbitrary number of omic modalities as input, for clarity we present them below for the simpler case of two omics; extension to more than 2 omics is straightforward. We denote the first omic as 𝑋^1^ and the second omic as 𝑋^2^, with 𝑁 denoting the number of samples, 𝑛^1^ the number of features (*e.g.* species) in 𝑋^1^, and 𝑛^2^ the number of features (*e.g.* metabolites) in 𝑋^2^. Each of the models can produce outputs 𝑋^⌃1^, representing the reconstruction or prediction of 𝑋^1^, and/or 𝑋^⌃2^, representing the reconstruction or prediction of 𝑋^2^. [𝑋^1^, 𝑋^2^] denotes a combined matrix where 𝑋^1^and 𝑋^2^are concatenated along the feature dimension. If 𝑋^1^ has dimensions 𝑁 × 𝑛^1^ and 𝑋^2^ has dimensions 𝑁 × 𝑛^2^ , the result if this concatenation is a matrix of dimensions 𝑁 × (𝑛^1^ + 𝑛^2^).

### The Diagonal (D) model

The Diagonal model employs a straightforward encoder-decoder architecture designed to predict one omic profile, from another, with an encoder that embeds the first omic into a lower dimensional latent space, and a decoder that predicts the second omic from this latent representation (Figure 1A).

More technically, the **encoder** maps high-dimensional input data 𝑋^1^ into a lower-dimensional latent representation 𝑍 . It consists of fully connected linear layers, where the number of hidden layers and the size of the latent layer are treated as hyperparameters. The number of neurons in the hidden layers decreases from the input to the latent. By default, each hidden layer has generally half as many units as the previous one (e.g., with two hidden layers and a 50-unit latent, a 400-unit input yields 400→200→100→50). When simple halving is not feasible (e.g., smaller inputs), we use evenly spaced widths between the input and the latent (e.g., 200→150→100→50) to maintain a smooth taper. To introduce non-linearity, each hidden layer is followed by a Rectified Linear Unit (ReLU) activation function. Formally: 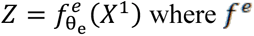 denotes the encoder function, and 𝜃_𝑒_ represents its learnable parameters.

The **decoder** predicts the second omic dataset, 𝑋^2^, from the latent space, 𝑍 . Its structure follows a design similar to the encoder, with the number of hidden layers treated as a hyperparameter and the number of neurons in each hidden layer defined using the same approach as in the encoder. ReLU activation functions are applied after each hidden layer to introduce non-linearity, except for the decoder’s final layer.

This process can be expressed mathematically as: 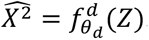

The final predictive model is the composition of the encoder and decoder:

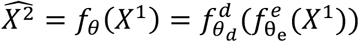

The model is trained to optimize parameter set 𝜃 = {𝑊_𝑒_, 𝑏_𝑒_, 𝑊_𝑑_, 𝑏_𝑑_} to minimize the following Mean Squared Error (MSE) loss function, which quantifies the difference between the true values of the second omic dataset 𝑋^2^, and its predicted values 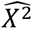:

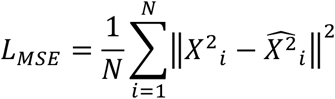

where 𝑁 represents the number of samples, 𝑖 denote a specific sample, 𝑋^2^ is the true second omic profile, and 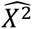 is the prediction of the second omic profile derived from the first omic profile 𝑋^1^.

### The Y model

The Y model extends the D model by training the encoder-decoder architecture to predict one omic from another while also reconstructing the first omic (Figure 1B).

Formally, the encoder and decoder function are defined as:

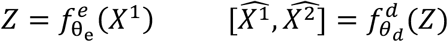

Where the decoder outputs a concatination of the reconstructed first omic 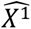 and the predicted second omic 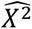.

The model is trained to minimize a combined MSE loss over both outputs:

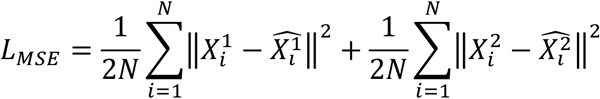

where 𝑁 represents the total number of samples, 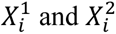 are the true values of the first and the second omic profiles for the 𝑖-th sample, and 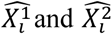 are their reconstructed and predicted values, respectively.

### The X model

The X model builds upon the Y model by alternating the input omic across training epochs, with the input being the first omic during even epochs and the second omic during odd epochs (Figure 1C). Specifically, the input is formed as [𝑋^1^, 0] during even epochs and [0, 𝑋^2^] during odd epochs. The encoder processes the input into a shared latent representation:

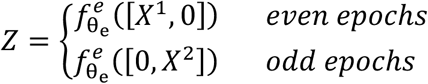

The decoder always reconstructs and predicts both omic, regardless of which was provided as input. The output is a full-length vector contraining both omic predictions:

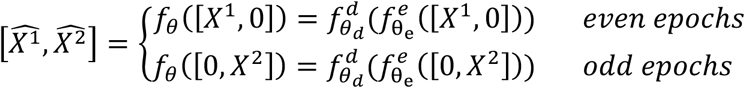

This design enables the model to reconstruct the input omic and predict the missing one. For example, when provided with [𝑋^1^, 0], the model reconstructs 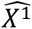 and predicts 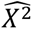; when given [0, 𝑋^2^], it reconstructs 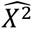 and predicts 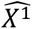. The full model is trained to minimize an MSE loss across both omics:

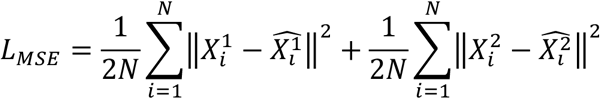

where 𝑁 represents the number of samples, 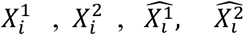 are the true and reconstructed/predicted values for sample 𝑖. This alternating structure allows the model to learn shared and omic/specific features while enabling bidirectional cross-omic prediction.

### The Parallel models (P)

The two parallel models, P_d_ and P_dp_, have similar architecture; they simultaneously train two autoencoders, each of the autoencoders corresponds to a different omic. The parallel model with the *inter-omic distance loss*, P_d_, ensures that samples close in one omic’s latent space are also close in the other. The parallel model with the *phenotype*-*distance loss*, P_dp_, further aims to align the latent space distances to distances between samples in the phenotype space. The following model architecture is relevant for the two models (P_d_ and P_dp_; Figure 1D).

The encoder of each omic maps the high dimensional input data 𝑋 (for X ∈ [X^1^, X^2^]) into a lower dimensional latent representation 𝑍. The network architecture is similar to the previous models. Formally:

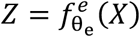

where 𝑓^𝑒^ is parameterized by 𝜃_𝑒_.

The decoder reconstructs the input data, *X* from the latent representation 𝑍, and mirrors the encoder structure:

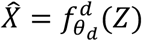

where 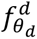 is another neural network parameterized by 𝜃_𝑑_.

The final predictive model is the composition of the encoder and decoder, mapping input data to its reconstruction via the latent space:

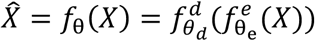

Specifically, both the P_d_ and P_dp_ models use reconstruction MSE loss which measures the difference between the original input 𝑋 and its reconstruction *X*:

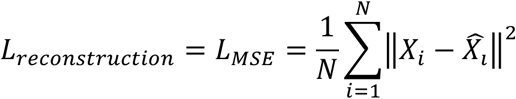

where 𝑁 is the number of samples, 𝑋_𝑖𝑘_ is the 𝑘-th feature for the original input sample 𝑖 and *X_ik_* is the 𝑘-th feature for the reconstructed output of sample 𝑖, for 𝑋 ∈ [𝑋^1^, 𝑋^2^].

Both models further incorporate an additional loss term (*inter-omic distance* loss), which is designed to ensure that samples close to one another in one omic’s latent space are also close to one another in the other. The combined loss function is thus of the form:

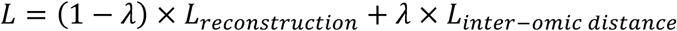

To calculate the distance loss, we calculate the pairwise Euclidean distance between all samples in each latent space. For each omic, this results in a Euclidean distance matrix 𝐷, where 𝐷_𝑖j_ = ||𝑍_𝑖_ − 𝑍_j_|| denotes the Euclidean distance between samples 𝑖 and 𝑗 in the latent space 𝑍. Notably, this calculation is feasible as the autoencoders for all omics are trained simultaneously. Next, we extract the upper triangular elements (not including the diagonal) from each distance matrix to focus on the unique pairwise distances, such that 𝑈 = {𝐷_𝑖j_|𝑖 < 𝑗}, where 𝑖 and 𝑗 are indices representing individual samples in the dataset. We calculate the Pearson correlation between the upper triangular vectors 𝑈 each omic’s distance matrices.

The *inter-omic distances loss* is finally calculates as:

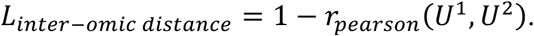

This loss component quantifies how well the latent spaces preserve distances between samples across omics.

λ, the weighting coefficient in the formulation about, acts as a regularization parameter that gradually increases during the training process:

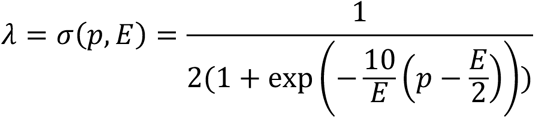

where 𝑝 is the epoch and 𝑁 is the total number of epochs and the factor 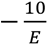 adjust the slope of the curve. This regularization initially assigns more weight to the reconstruction loss and subsequently to the distance loss, ensuring a balanced optimization.

**The P_dp_ model** is similar to the P_d_ model, but also ensure that distances in the latent spaces align with distances in the phenotype space. In this model, the overall loss function is defined as:

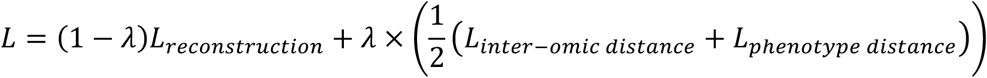

The calculation of the 𝐿_𝑝ℎ𝑛𝑜𝑡𝑦𝑝𝑒 𝑑𝑖𝑠𝑡𝑎𝑛𝑐𝑒_ is similar to that of the 𝐿_𝑖𝑛𝑡𝑒𝑟–𝑜𝑚𝑖𝑐 𝑑𝑖𝑠𝑡𝑎𝑛𝑐𝑒_. Specifically, we first calculate the distance matrix based on the categorical or continuous phenotype of each sample (𝐷^𝑝^) and similarly extract the upper triangular elements to focus only on the unique pairwise distances, excluding redundant entries and the diagonal 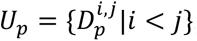, where 𝑖 and 𝑗 are indices representing individual samples in the dataset. We again calculate the Pearson correlation between the upper triangular of distance matrices for each of the specific omic and of the phenotype. For example, for the first omic, the *phenotype distances loss* is calculated as follows:

The same calculation is performed for the second omic, yielding:

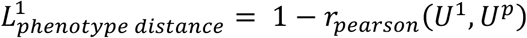

These correlations quantify how well the latent spaces distances align with phenotypic distances.

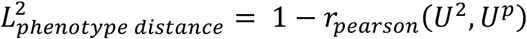

### Training Procedure (for all models)

Each model is trained using 5-fold cross-validation, where the dataset is randomly split into five equal folds. In each iteration, the model is trained on four folds and validated on the remaining one. The final performance is averaged across all folds. We use the Adam optimizer, with the learning rate treated as a hyperparameter. The learning rate is selected through hyperparameter tuning, and it is typically initialized at 0.0001 or 0.001. To prevent overfitting, we further apply weight decay (L2 regularization) with a value of 1 × 10^−5^, which penalizes large weights and promotes better generalization. We set the Adam optimizer’s ε (epsilon) parameter to 1 × 10^−8^, a small constant in the denominator of the update rule that prevents division by zero and improves numerical stability without affecting the optimization objective. We trained all models with a latent layer size of 32 or 64 neurons and without additional hidden layers beyond the latent representation. The training process is performed using mini-batch gradient descent, with a batch size of 16 for the Farnzosa *et al.* dataset and 32 for the Lifelines and the DAP datasets. Models were trained for either 500 or 1,000 training epochs. Both the latent layer size and the number of epochs can be selected via hyperparameter tuning. All models are implemented using PyTorch (version 2.1.2), and are designed to run on both CPU and GPU, with GPU acceleration supported via CUDA for faster training when available.

To enable a direct and fair comparison across model architectures, we kept the hyperparameter configuration fixed across all models. For the parallel architectures, we also applied the same hyperparameters to both subnetworks. While independently tuning each network or model type could further improve absolute performance, we intentionally maintained consistent hyperparameters to ensure that performance differences reflected the model architecture rather than differences in tuning.

### Models evaluation

We evaluate each model on two tasks: (i) cross-omic prediction (predicting the metabolome from the microbiome) and (ii) phenotype prediction from test-set latent embeddings. These assess how well models capture cross-omic structure and phenotype-relevant signal.

For each metabolite, accuracy was measured as the Spearman correlation between observed and predicted values. A metabolite is well-predicted if r ≥ 0.3 and FDR < 0.1; we report the mean count of well-predicted metabolites across folds. In D, Y, X the metabolite predictions come directly from the model output. In P_d_ and P_dp_, we fit a linear map in latent space on the training fold, 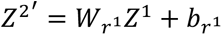, then decode to the metabolome 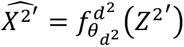. The regression is implemented using the LinearRegression function from the scikit-learn library, (version 1.3.2).

A Random Forest (RF) regressor was trained on the training latent embeddings to predict a continuous phenotype. Test samples were encoded with the pretrained encoder and scored by the trained RF. Accuracy is reported as the mean Spearman correlation between observed and predicted values across folds. For model evaluation, D and Y provide a single latent space; X is a single model evaluated with either omic alone; P_d_ and P_dp_ include two encoders (two latent spaces), and we report prediction using either omic as input. The RF is implemented using the RandomForestRegressor function from the scikit-learn library, (version 1.3.2), using 100 estimators and default hyperparameters. Models were fit in each fold on the training partition and evaluated on held-out test data.

### Mantel test for microbiome-metabolome association

We quantified the global association between microbiome and metabolome profiles using the Mantel test. Specifically, for each cohort, we built sample-matched distance matrices: Bray-Curtis dissimilarities from microbiome relative-abundance tables, and Euclidean distances from metabolite profiles. We then applied a Pearson Mantel test with 999 permutations (scikit-bio, version 0.5.8), reporting the Mantel correlation and permutation p-value.

### Evaluation of association of the latent space and input data with the phenotype

We quantified the association between latent-space geometry and the phenotype using PERMANOVA (adonis2, vegan v2.6-4). For each analysis, we computed pairwise Euclidean distances among samples from the embeddings (or the input features, where noted) and tested whether the phenotype explained variation in these distances using 9,999 permutations.

## Discussion

In this study, we presented and evaluated five encoder-decoder models designed to integrate multi-omic data, exploring their ability to support both cross-omic prediction and improved phenotype prediction. The various analyses presented above allowed us to systematically characterize how latent representations that are derived from multi-omic data capture key relationships within and across omic layers and how different architectures impact this capacity.

Of the models we studied, the three ‘shared-space’ encoder-decoder models (D, Y, and X) prioritize simplicity and apply straightforward architectures for predicting one omic from another, making them easily accessible for diverse applications. The two ‘parallel’ multi-autoencoder models (P_d_, and P_dp_), in contrast, utilize a fundamentally different approach with multiple latent spaces, one per each omic, to enhance the flexibility of these models. Specifically, this design enables generating independent, yet interconnected representations of the various omics, thus improving predictive performances and offering insights into the cross-omic associtions.

As noted above, these five models represent a progression in network architecture, in how multi-omic data are integrated, and in the way underlying data structures are being captured. We began with the D model, a simple baseline encoder-decoder architecture for predicting one omic from another. The Y model extended this by also reconstructing the input omic, forcing the latent space to retain also omic-specific information about the input omic. The X model further generalized this approach by alternating between omics during training to reconstruct and predict each from the other, potentially allowing the model to learn both shared and distinct patterns. Moving beyond single-network designs, the P_d_ and P_dp_ models then utilized multiple parallel autoencoders, each dedicated to a different omic, with separate latent spaces. The P_d_ model aligns these latent spaces to each other by forcing them to incorporate a similar pairwise distance structure between samples, while the P_dp_ model additionally incorporates phenotype distances into the alignment, promoting biologically relevant embeddings.

We evaluated the models on two key tasks: predicting one omic based on the other, and predicting a continuous phenotype from the learned latent representations. These tasks were assessed on three independent datasets: a relatively small IBD cohort with fecal microbiome and fecal metabolome (and calprotectin levels as a phenotype of interest), and two larger datasets combining fecal microbiome and serum metabolome (with age as the phenotype). In the cross-omic task, we found that the Y and X models consistently improved upon the D model, likely by capturing richer latent representations. The parallel models, P_d_ and P_dp_, further improved upon the three simpler models, suggesting that the ability to construct parallel latent spaces supports a more expressive and modular foundation. In phenotype prediction, all models exhibited comparable performances in predicting the phenotype using a microbiome profile as input. However, the parallel models, as well as the X model, further supported predicting the phenotype based on metabolome profile input, exhibiting markedly improved prediction, with the P_dp_ model, perhaps not surprisingly, achieving the best results. Interestingly, the X model also achieved competitive results, highlighting the value of cross-omic training even with a single latent space.

Importantly, this ability of the parallel models to generate distinct latent representations for each omic has several advantages. First, it enables independent phenotype predictions from each data type and a direct comparison of the predictive power of each omic. This could potentially further facilitate ensemble prediction models that integrate predictions from the various omics. Such comparisons not only help identify which omic is more closely tied to the phenotype of interest but may also reveal omic-specific biomarkers, latent structures, or discrepancies that point to confounding factors, hidden subtypes, or novel biological signals detectable in only one omic layer. More generally, such models offer an independent *embedding* for each omic, which could be beneficial for multiple downstream algorithms and methods, for example, for multi-view clustering (subtyping), integrative network analysis, or causal inference (e.g., mediation analysis) to determine how one omic layer influences another in a biological pathway.

It should also be noted that unlike the D and Y models, the X, P_d_, and P_dP_ architectures do not require a pre-specified directionality between omics, that is, they do not rely on defining one omic as input and the other as output. In the study above, we used the microbiome as input to predict the metabolome, as microbial composition is often considered a driver of metabolic profiles^60^. Accordingly, phenotype prediction results for the D and Y models are reported above only using the microbiome as input. For the sake of completeness, however, we also evaluated the reverse direction, using the metabolome as input to predict the microbiome, and found that the overall trends remained consistent, with the P_dP_ model still outperforming the alternatives (Supplementary Figure 10).

We believe that the architectural progression presented in our study, from single encoder-decoder models to parallel, phenotype-aligned ones, provides an intuitive framework for understanding how architectural enhancemant can improve predictive performance. This stepwise evolution of architectures clarifies the rationale behind each model, demystifying the use of neural networks in multi-omic research, and making these tools more accessible for broader applications. The P_dp_ model’s success in aligning the latent space to a continuous phenotype also opens the door to future extensions involving more complex or multi-dimensional phenotypes.

Despite encouraging results, however, several limitations warrant attention. For example, the first dataset analyzed was relatively small for multi-omic modeling with continuous phenotypes. While larger datasets were used for generalization, they posed other challenges, such as weaker microbiome-metabolome associations and a potentially more challenging phenotype. Moreover, incorporating additional omics, such as metatranscriptomics or epigenomics, could enrich the latent representations and improve both prediction and biological insight. However, such datasets remain limited in availability, and their integration poses challenges due to differences in scale, dimensionality, and data structure. Notably, we believe that the parallel models, which allow each omic to be embedded into a separate latent space with potentially different regularities, would naturally cope with such differences in omic characteristics better than models that use a shared latent space and that force all omics to be mapped to this shared space despite their different characteristics. Our models also did not explicitly control for confounding factors (e.g., diet, demographics, or technical variability), which may influence associations. Addressing these confounders typically requires access to well-annotated metadata and can be done by incorporating them directly into the model. Lastly, while we used several sets of hyperparameters to fairly compare architectures, optimal performance would require tuning each model individually. Nonetheless, the performance trends we observed were consistent across different hyperparameter configurations, suggesting that the results reflect general model behavior rather than specific parameter choices.

Overall, this study introduces a suite of network architectures designed to integrate multi-omic data. By systematically building model complexity, from single latent spaces to parallel representations aligned with phenotypic structure, we show how architectural choices impact both prediction accuracy and the biological resolution of latent representations, offering robust, systematic, and flexible foundations for integrative multi-omic analysis.

## Code availability

The python code used for this analysis includeing the PAPRICA framework is available on GitHub (https://github.com/borenstein-lab/PAPRICA).

## Acknowledgements

We thank members of the Borenstein lab for helpful advice and discussions.

This study was supporting in part by NIH U19 grant AG057377, and by a research grant from the Center for AI & Data Science (TAD) at Tel Aviv University. T.B. was supported in part by a fellowship from the Edmond J. Safra Center for Bioinformatics at Tel-Aviv University.

The Lifelines initiative has been made possible by subsidy from the Dutch Ministry of Health, Welfare and Sport, the Dutch Ministry of Economic Affairs, the University Medical Center Groningen (UMCG), University of Groningen and the Provinces in the North of the Netherlands (Drenthe, Friesland, Groningen).

This research is based on publicly available data collected by the Dog Aging Project, under U19 grant AG057377 (PI: Daniel Promislow) from the National Institute on Aging, a part of the National Institutes of Health, and by additional grants and private donations, including generous support from the Glenn Foundation for Medical Research, the Tiny Foundation Fund at Myriad Canada, and the WoodNext Foundation. These data are housed on the Terra platform at the Broad Institute of MIT and Harvard. The authors thank Dog Aging Project participants, their dogs, and community veterinarians, as well as all Dog Aging Project Team members for their important contributions.

The icons used in Fig. 1 were designed by ‘Freepik’ and are available through the ‘Flaticon license’ (www.flaticon.com).

## Dog Aging Project Consortium

Joshua M. Akey | Rozalyn M. Anderson | Elhanan Borenstein | Marta G. Castelhano | Amanda E. Coleman | Kate E. Creevy | Matthew D. Dunbar | Virginia R. Fajt | Jessica M. Hoffman | Erica C Jonlin | Matt Kaeberlein | Elinor K. Karlsson | Kathleen F. Kerr | Jing Ma | Evan L. MacLean | Stephanie McGrath | Natasha J Olby | Daniel E.L. Promislow | May J Reed | Audrey Ruple | Stephen M. Schwartz | Sandi Shrager | Noah Snyder-Mackler | M. Katherine Tolbert

## Supplementary information

**Supplementary Figure 1:**
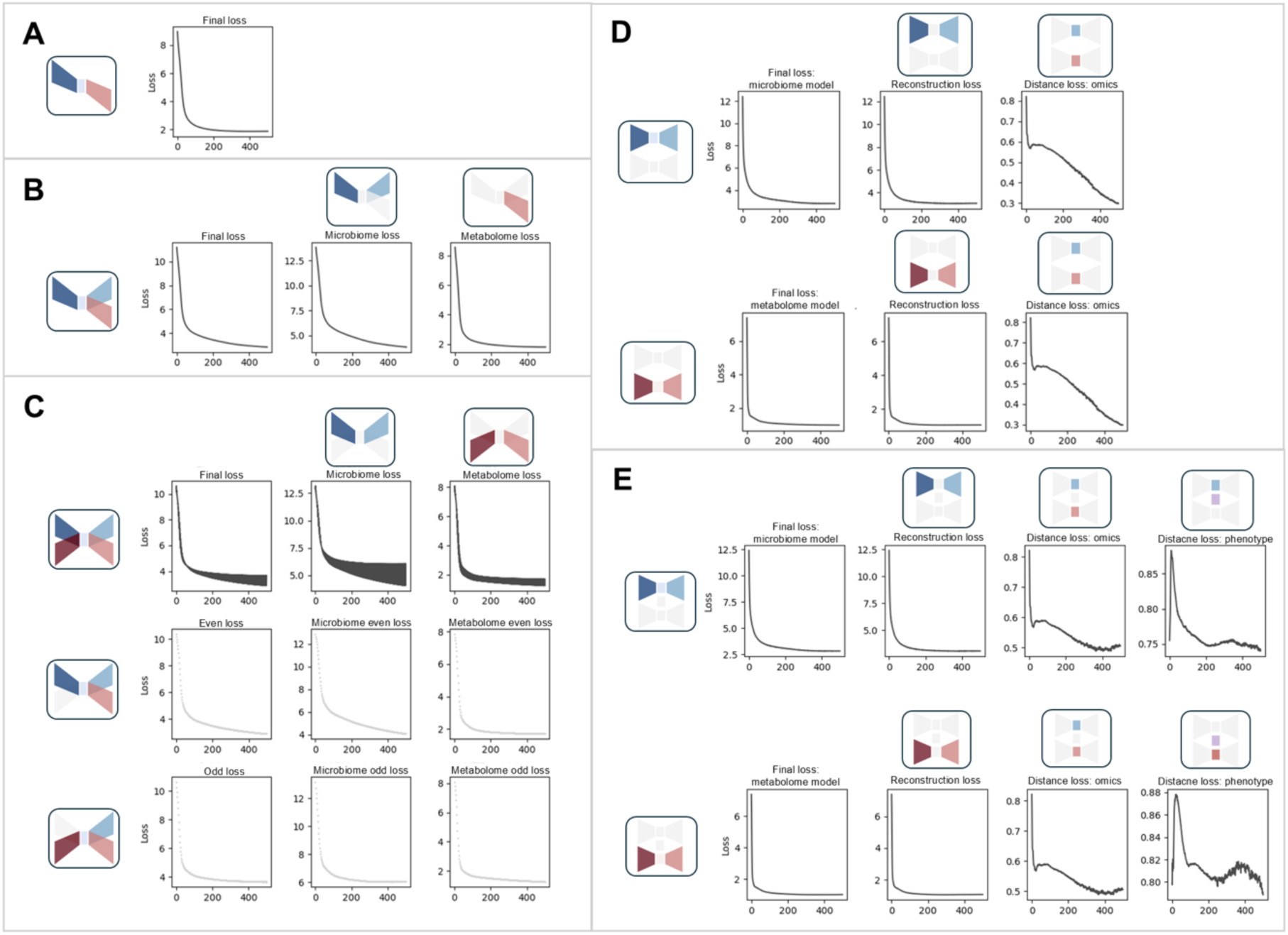
Training dynamics and convergence patterns of the proposed models on *unseen test* data from the Franzosa *et al*. microbiome-metabolome dataset. **A.** Loss curves for the D model, representing the Mean Squared Error (MSE) loss between the real metabolome data and the predicted metabolome data generated from microbiome inputs. **B.** Loss dynamics for the Y model, showing from left to right: (i) the final loss, comprising two MSE loss components; (ii) MSE loss between the input microbiome data and the reconstructed microbiome; and (iii) MSE loss between the real metabolome data and the predicted metabolome data. **C.** Loss dynamics for the X model with the top row showing the overall loss across all alternating inputs, the second row showing the loss when microbiome data is used as input (even epochs), and the third row showing the loss when metabolome data is used as input (odd epochs). In each row, the panels (from left to right) show: (i) the final loss, composed of two MSE loss components; (ii) MSE loss between the input microbiome data and the reconstructed microbiome; and (iii) MSE loss between the real metabolome data and the reconstructed metabolome data. **D.** Loss curves for the P_d_ model, where the upper row represents the microbiome model and the lower row represents the metabolome model, both trained simultaneously. Panels (from left to right) show: (i) the final loss, which includes reconstruction and *inter-omic distance losses*; (ii) reconstruction loss between input and output data (microbiome in the upper row, metabolome in the lower row); (iii) the *inter-omic distance loss*, which quantifies distances between omics in the latent space (microbiome in the upper row, metabolome in the lower row). **E.** Loss curves for the P_dp_ model, with the same layout and loss components as in D, but including the second component of the distance loss: (iv) *phenotype-distance loss*, which aligns latent space distances (microbiome in the upper row, metabolome in the lower row) with phenotype distances.

**Supplementary Figure 2:**
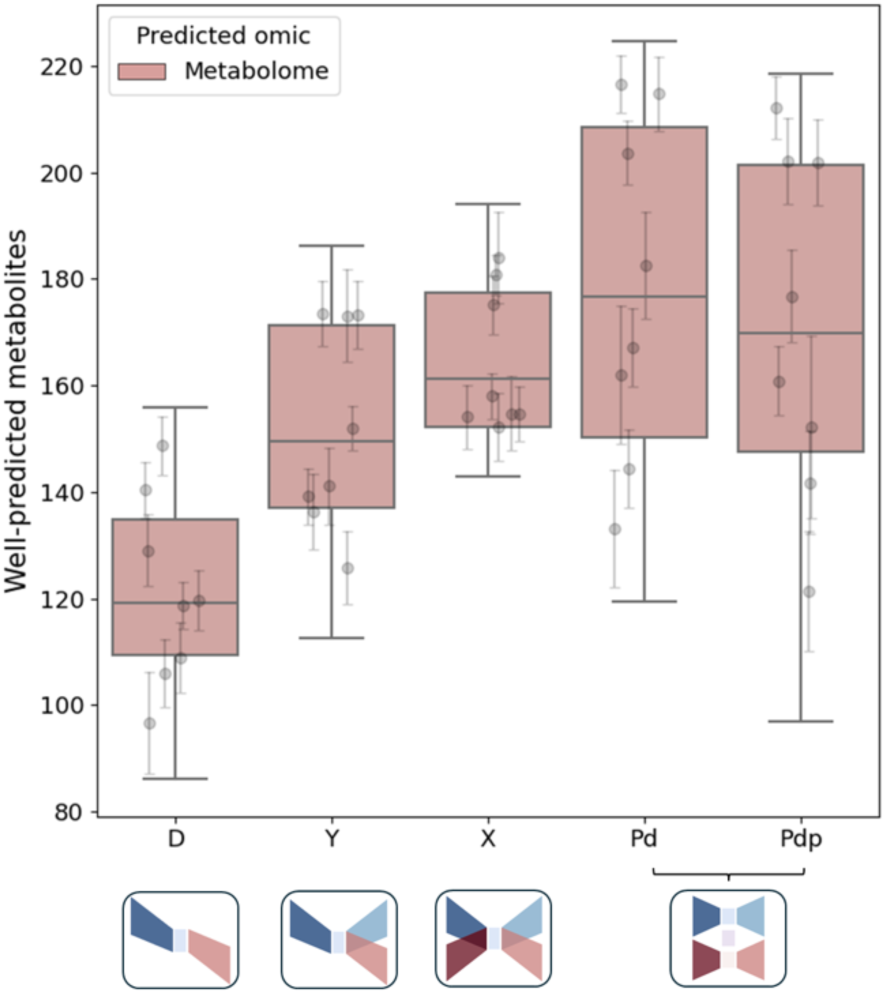
Stability of metabolite prediction across model initializations. Box plot showing the number of well-predicted metabolites (Spearman’s r > 0.3, FDR < 0.1) for each of the five models across different hyperparameter settings. Each dot represents a single run with a specific hyperparameter configuration, while error bars indicate the standard deviation of the number of well-predicted metabolites across multiple runs of the same model with identical hyperparameters but different random initial weights.

**Supplementary Figure 3:**
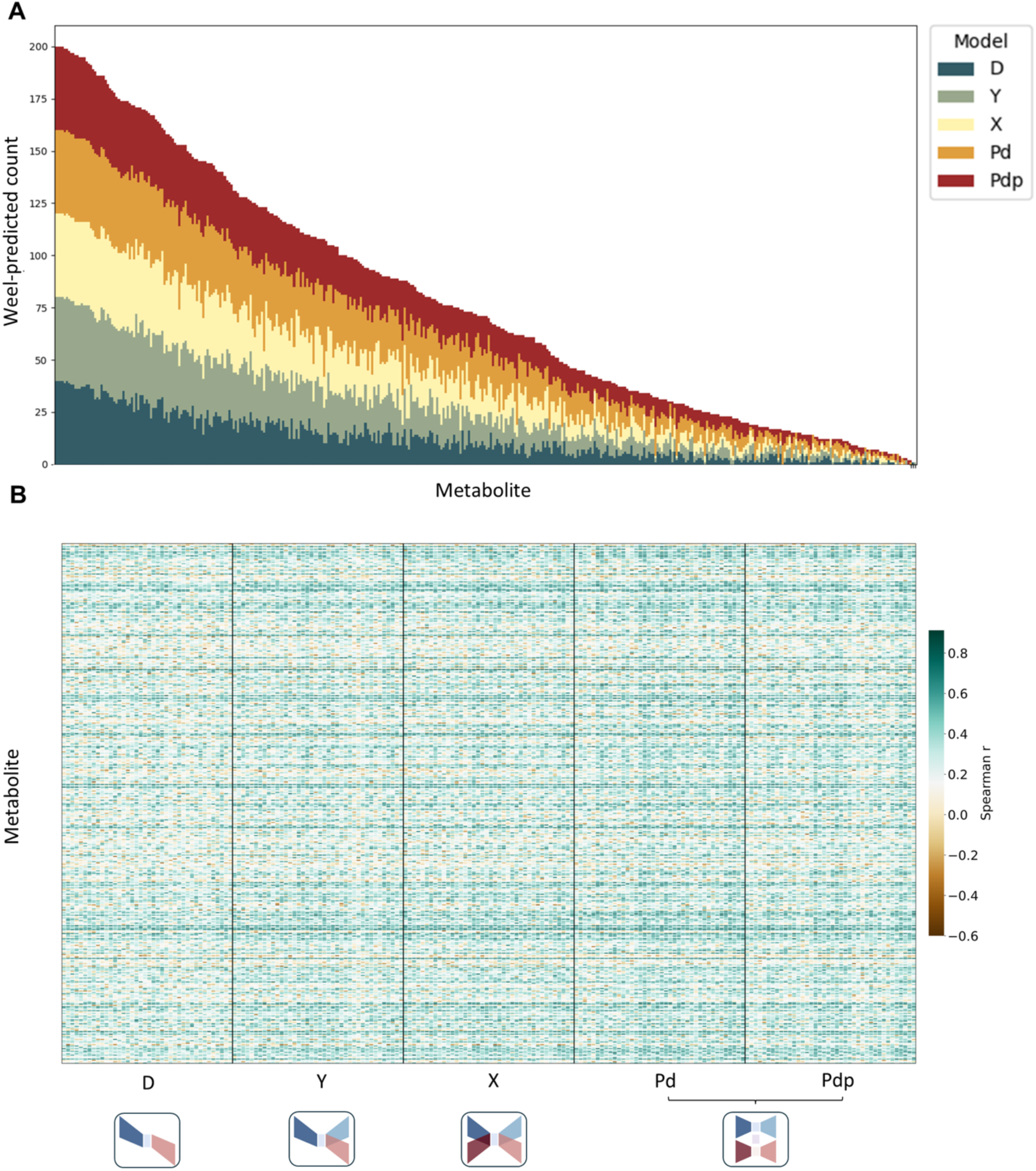
Consistency of metabolite prediction across hyperparameters and cross-validation folds on Franzosa et al. dataset. **A.** Stacked bar plot showing the number of times each metabolite was classified as well predicted across different hyperparameter settings and cross-validation folds. Only metabolites that were observed as well predicted at least once are included. Each column represents a metabolite, and the height of the bar indicates its prediction consistency. The stacked segments within each bar represent different models, with each color corresponding to a specific model.**B.** Heatmap displaying the correlation between observed and predicted metabolite values (computed across samples within each run). Rows representing metabolites and columns representing runs (i.e., different hyperparameter settings and cross-validation folds), ordered by model. Color intensity reflects the correlation strength.

**Supplementary Figure 4:**
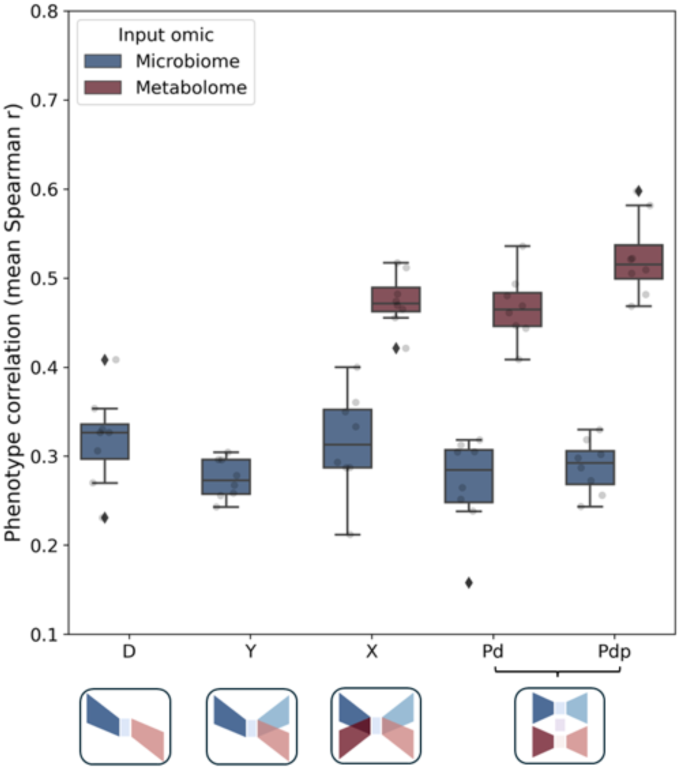
Evaluation of model performance in phenotype prediction using linear regression. Same as Figure 3C, but here phenotype prediction was performed using a linear regression model rather than a random forest model.

**Supplementary Figure 5:**
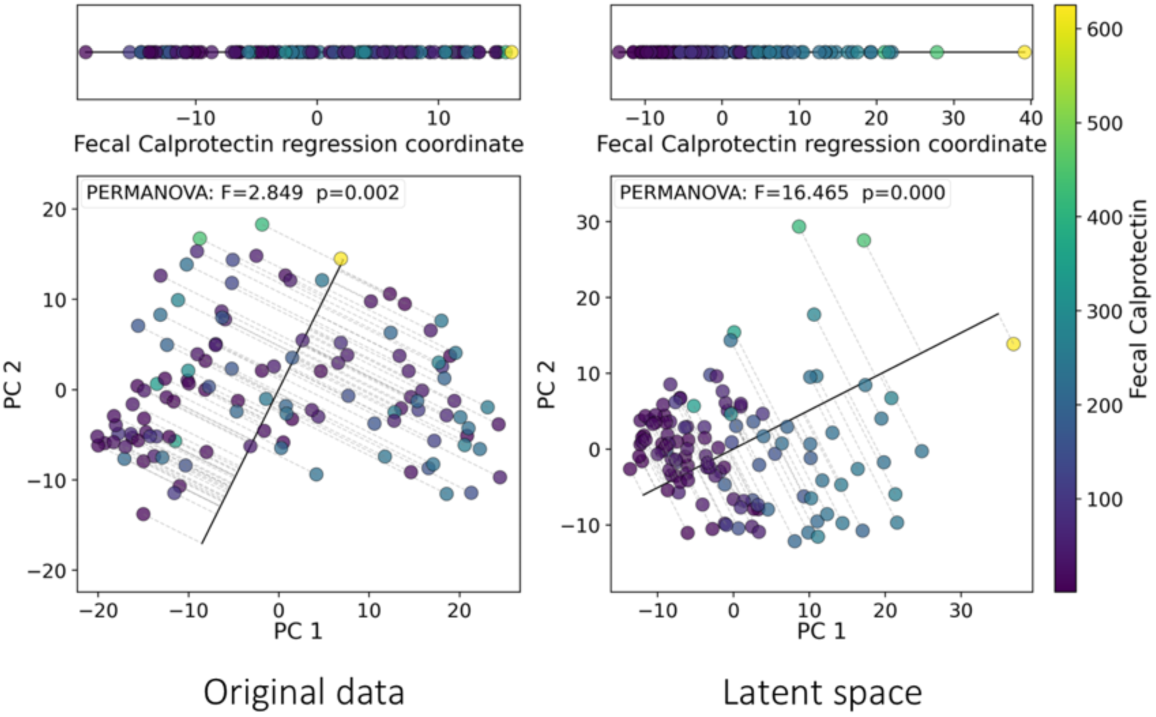
Principal Component Analysis (PCA) of the metabolome data and its latent space representation in the training set. Same as Figure 2D, but performed using the training data.

**Supplementary Figure 6:**
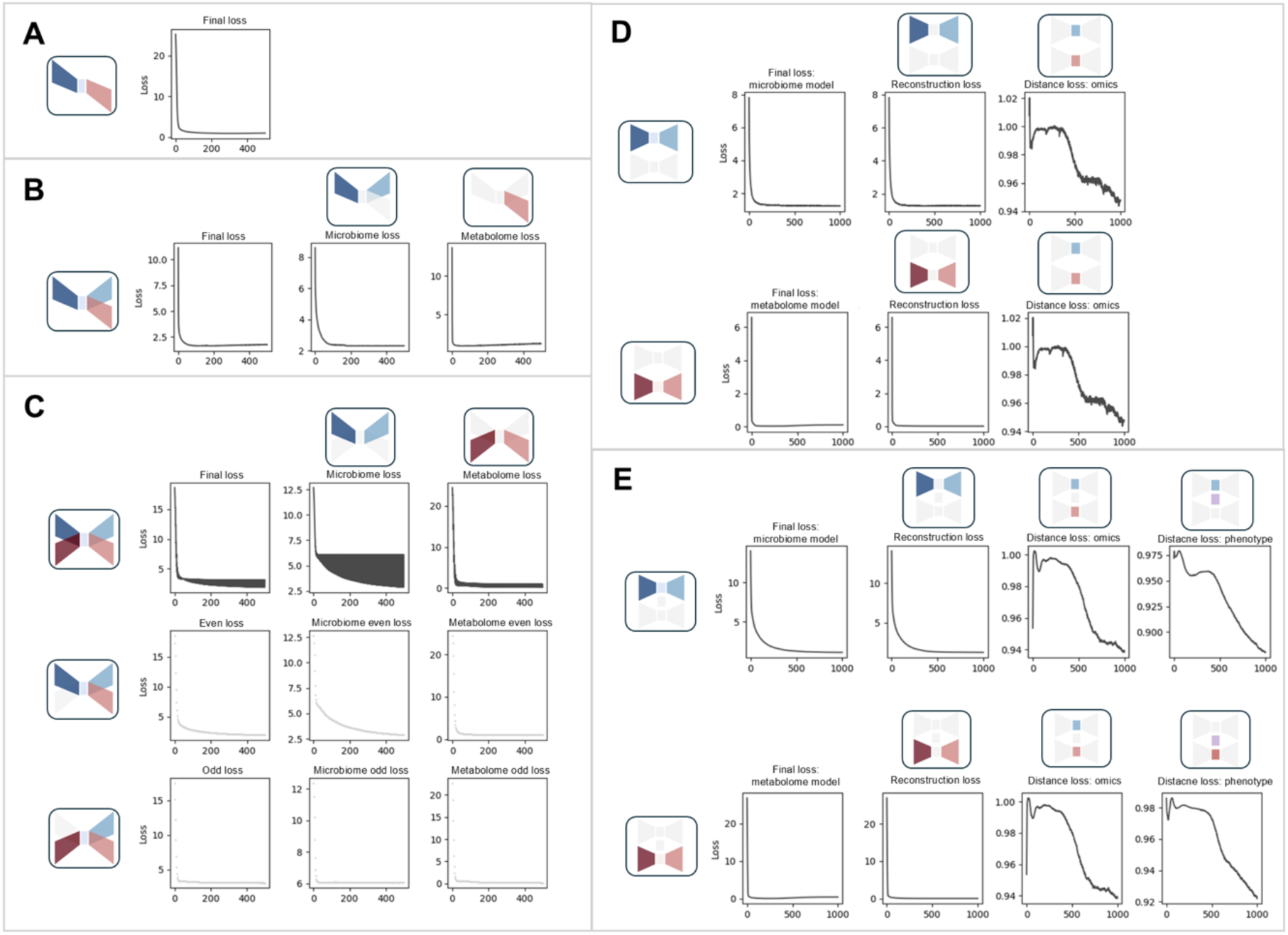
Training dynamics and convergence patterns of the proposed models on test data, using the Lifelines dataset. **A-E.** Same as Supplementary Figure 1A-E, but using the Lifelines dataset.

**Supplementary Figure 7:**
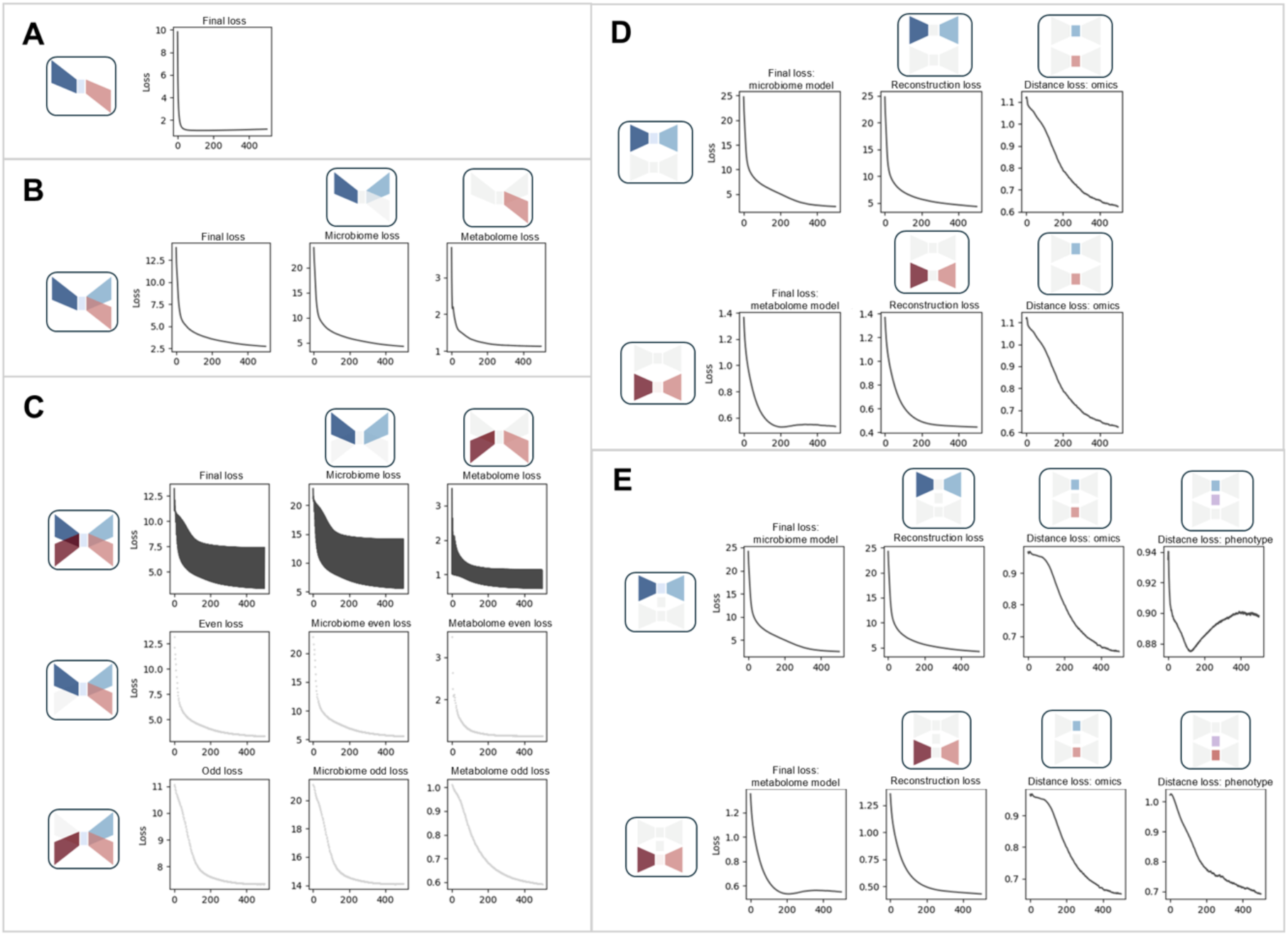
Training dynamics and convergence patterns of the proposed models on test data, using the Dog Aging Project (DAP) data. **A-E.** Same as Supplementary Figure 1A-E, but using the DAP dataset.

**Supplementary Figure 8:**
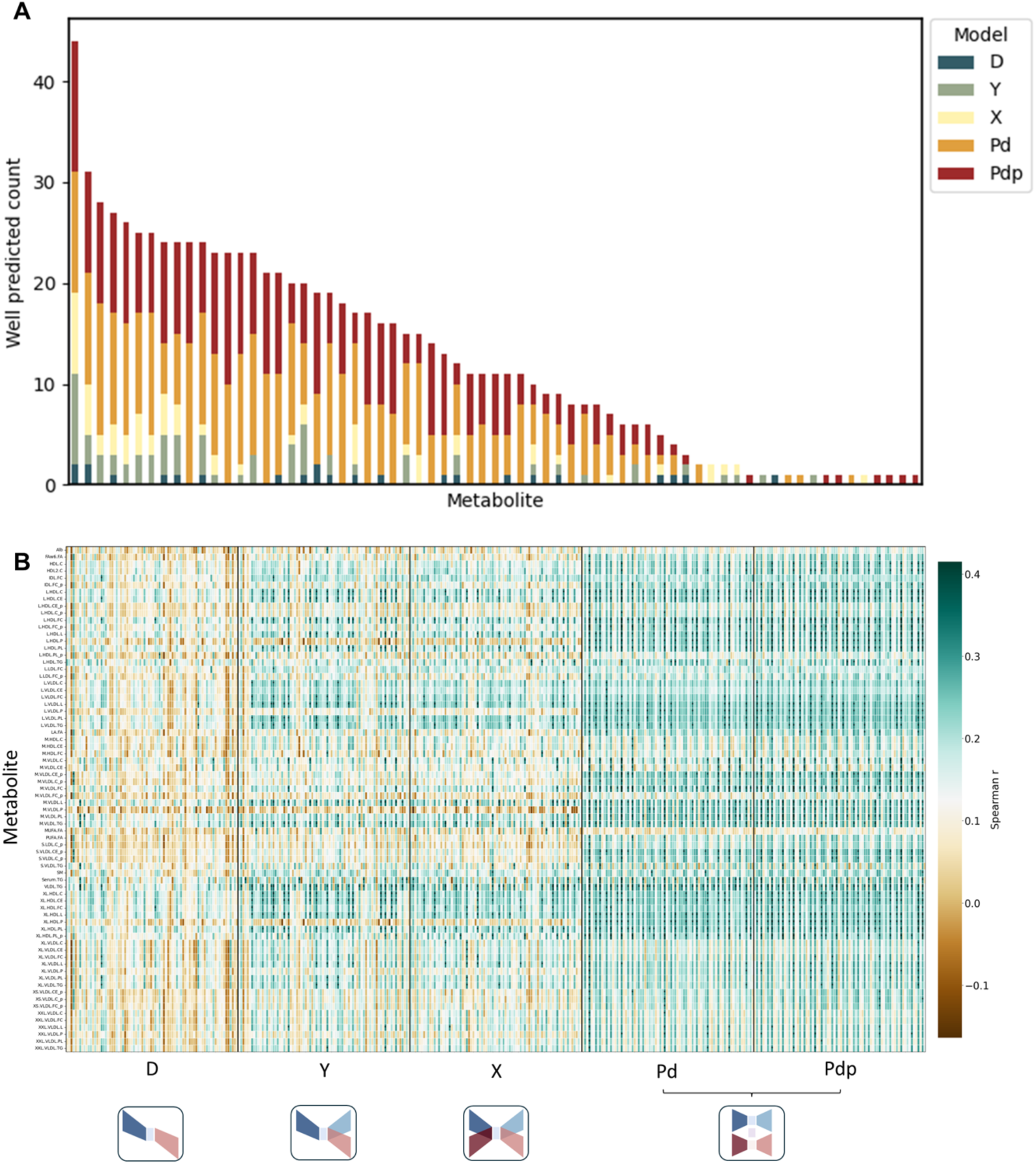
Consistency of metabolite prediction across hyperparameters and cross-validation folds on Lifelines dataset. **A.** Same as Supplementary Figure 3A, but using the Lifelines dataset. **B.** Same as Supplementary Figure 3B, but using the Lifelines dataset. Only metabolites that were observed as well predicted at least once are included. Color intensity reflects the correlation strength. Well-predicted metabolites in specific runs are marked with asterisks.

**Supplementary Figure 9:**
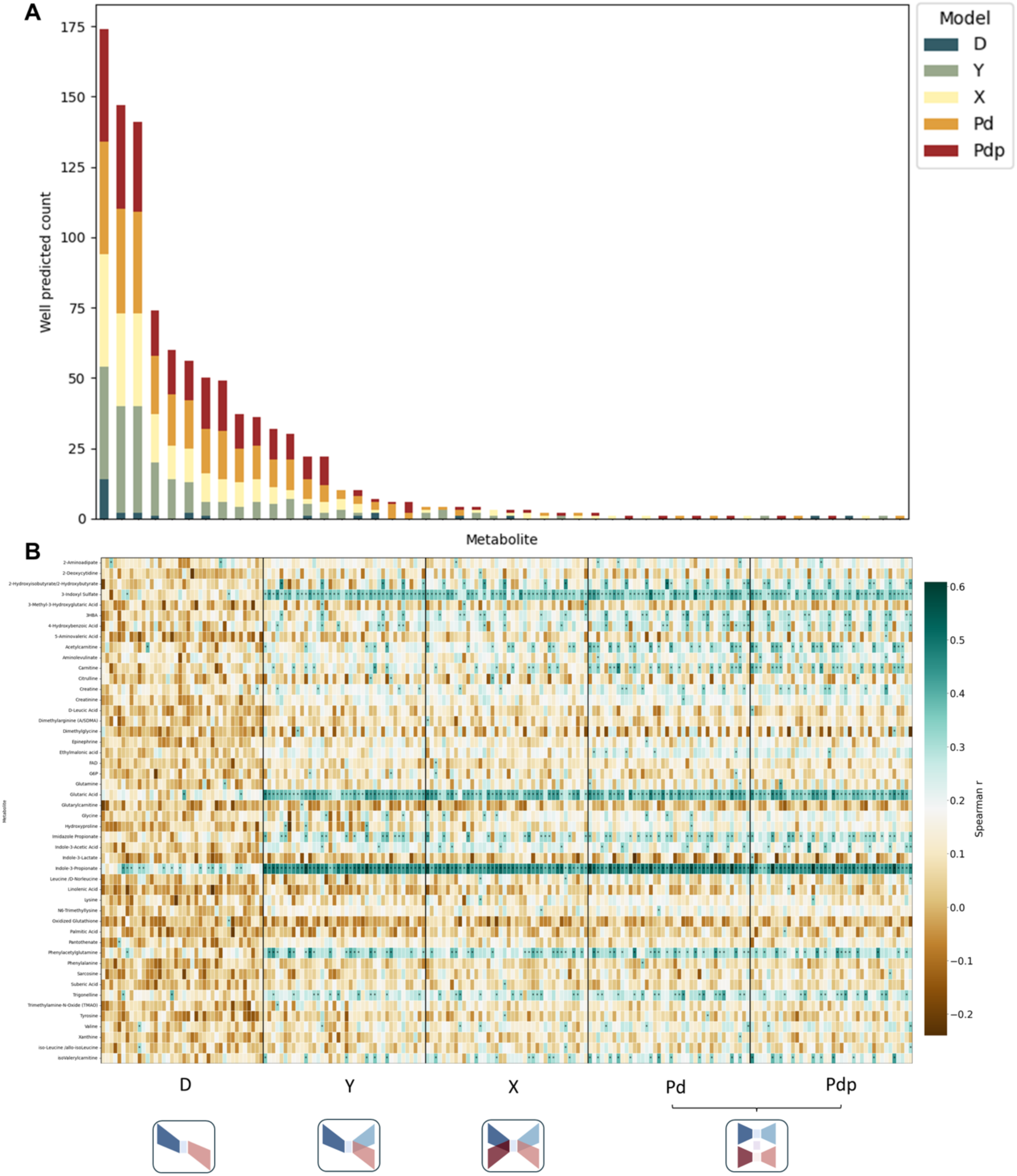
Consistency of metabolite prediction across hyperparameters and cross-validation folds on the DAP dataset. **A.** Same as Supplementary Figure 3A, but using the DAP dataset. **B.** Same as Supplementary Figure 3B, but using the DAP dataset. Only metabolites that were observed as well predicted at least once are included. Color intensity reflects the correlation strength. Well-predicted metabolites in specific runs are marked with asterisks.

**Supplementary Figure 10:**
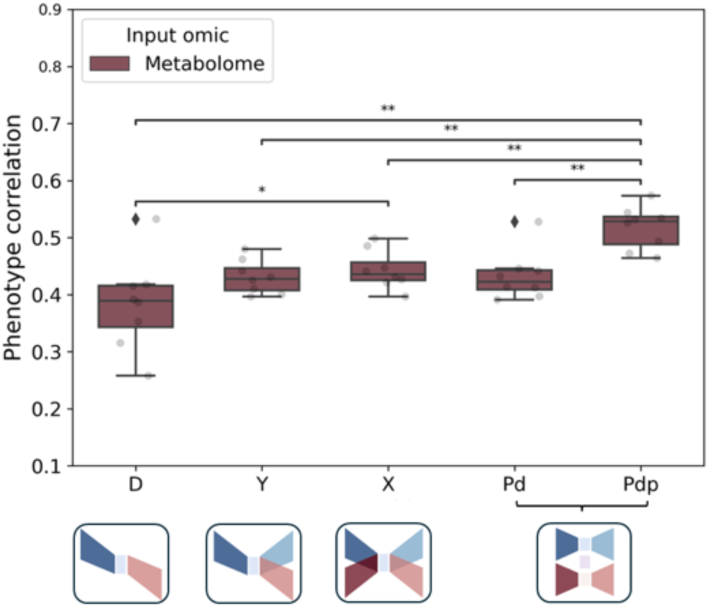
Evaluation of model performance in phenotype prediction using metabolome data. Same as Figure 3C, but here models’ performance is shown when using only the metabolome omic as input.

